# A deep learning network for parallel self-denoising and segmentation in visible light optical coherence tomography of human retina

**DOI:** 10.1101/2022.11.25.518000

**Authors:** Tianyi Ye, Jingyu Wang, Ji Yi

## Abstract

Visible light optical coherence tomography (VIS-OCT) of human retina is an emerging imaging modality that uses shorter wavelength in visible light range than conventional near infrared (NIR) light. It provides one-micron level axial resolution to better separate stratified retinal layers, as well as microvascular oximetry. However, due to the practical limitation of laser safety and comfort, the permissible illumination power is much lower than NIR OCT which can be challenging to obtain high quality VIS-OCT images and subsequent image analysis. Therefore, improving VIS-OCT image quality by denoising is an essential step in the overall workflow in VIS-OCT clinical applications. In this paper, we provide the first VIS-OCT retinal image dataset from normal eyes, including retinal layer annotation and “noisy-clean” image pairs. We propose an efficient co-learning deep learning framework for parallel self-denoising and segmentation simultaneously. Both tasks synergize within the same network and improve each other’s performance. The significant improvement of segmentation (2% higher Dice coefficient compared to segmentation-only process) for ganglion cell layer (GCL), inner plexiform layer (IPL) and inner nuclear layer (INL) is observed when available annotation drops to 25%, suggesting an annotation-efficient training. We also showed that the denoising model trained on our dataset generalizes well for a different scanning protocol.

## 1. Introduction

Optical Coherence Tomography (OCT) is a widely used imaging technique in ophthalmology to evaluate anatomical layers in the retina for diagnosis [1]. Current commercialized OCT devices use near-infrared (NIR) light source, either at around 850nm by spectral domain OCT or 1050nm by swept source OCT [2]. The recent emerging visible light optical coherence tomography (VIS-OCT) uses a shorter wavelength with visible light centered around 550nm for its illumination [3]. The resulting advantage is higher imaging contrast and one-micron level resolution, which allows more accurate analysis for 2D/3D retinal layers in both clinical applications as well as preclinical animal models [4–7]. Another advantage of VIS-OCT is its spatio-spectral analysis within the microvasculature for label-free oximetry (*i*.*e*., measuring hemoglobin oxygen saturation, sO_2_) [8–10]. The 3D imaging capability provides accurate vessel lumen segmentation that allows isolating signal within microvasculature to avoid other confounding signals as in other fundus-based oximetry. By that, microvascular retinal oximetry down to capillary level has been reported by VIS-OCT [11–13]. Clinical feasibility of microvascular sO_2_ in parafoveal vessels that are around 20-30 μm in diameter have been demonstrated in several retinal vascular pathologies [14,15]. In addition, due to the distinct scattering contrast in VIS-OCT in comparison with NIR-OCT, spectroscopic analysis can provide structural properties beyond image resolution. Song *et al*. demonstrates VIS-OCT reflectivity and spectroscopy of peripapillary retinal nerve fiber layer (pRNFL) better differentiate normal eyes with pre-perimetric eyes, implying an early detection method for glaucoma [16,17]. Gupta *et al*. utilized VIS-OCT with energy concentrated at discrete red, green, and blue wavelength bands to quantify macular pigments and localize them in depth within the human retina in vivo [18]. The spectral contrast provided by VIS-OCT can also be used for single-scan OCTA [19].

While the benefits of VIS-OCT in retinal imaging are clearly presented in preclinical and clinical studies, the challenge is the more stringent illumination power limit (∼0.1-0.25 mW at eye pupil) in VIS-OCT than in NIR OCT (∼1-2 mW). The shorter wavelength also makes VIS-OCT more susceptible to poor optical quality in aging and pathological eyes. Both will lead to degradation of image quality, compromising the ability to perform accurate segmentation, a necessary step for essentially all VIS-OCT quantitative analysis. Therefore, denoising VIS-OCT image and efficient/accurate layer segmentation become critical steps in the general workflow in VIS-OCT.

Recently, several supervised deep learning (DL)-based denoising methods [20,21] have been proposed, with the need for paired clean images as ground truth. At the same time, automatic segmentation of retinal layers has been intensively studied using DL, outperforming traditional graphical and machine learning (ML) methods [22–26]. However, most of these methods require a large amount of manual annotation by clinicians for training. While many DL approaches have been shown to be successful in each task, considering the real-world clinical scenarios, in which both denoising and segmentation are necessary, an efficient way to solve both the problems simultaneously is a significant need.

In this paper, we reported an efficient DL method for simultaneous denoising and segmentation for high-resolution VIS-OCT images. We collected and published the first VIS-OCT dataset with “noisy-clean” image pairs and ten manually delineated retinal boundaries. Inspired by DenoiSeg [27], we proposed a co-learning framework based on residual-UNet for simultaneous denoising and segmentation (named DenoiSegOCT). Different from DenoiSeg [27], our approach extends the self-supervised strategy Noise-2-Void(N2V) [28] for inherent noise reduction; and extends the 3-class segmentation (background, foreground, and edges) to 10-class segmentation. For comparison, we also provided supervised denoising strategy Noise-2-Label(N2L) in DenoiSegOCT. Such co-learning process significantly reduces the hyperparameters tuning time and simultaneously provide both denoised image and prediction of segmentation [27]. The experimental results suggested that the N2V denoising process helped the segmentation when available annotation drops to 25%. And the self-supervised denoising performance of our framework was qualitatively better than using N2V alone. Our model also generalized well for two different scanning protocols, indicating the robustness of our framework.

## 2. Methods

### 2.1. VIS-OCT dataset

This paper presents the first VIS-OCT human retina image dataset for machine-learning research. The data include retinal B-scans acquired by our 2^nd^ Gen dual-channel VIS-OCT system. The technical description of the device has been detailed previously [29]. Briefly, VIS-OCT covers a bandwidth of 500-640 nm by a linear-in-K spectrometer. The axial resolution is up to 1.3 um in tissue. The A-line rate is 100 kHz. The power in VIS-OCT is 0.2-0.24 mW on cornea. The device also implemented per A-line noise cancellation method to achieve near shot-noise imaging performance. The image processing methods are described in detail in [29]. Briefly, the reference arm spectra were recorded for each A-line from a second spectrometer and subtracted from the raw data to remove excessive noise. The image formation includes a digital dispersion compensation, and Fourier transform to generate B-scan images.

The scanning protocol (HD protocol) used a speckle reduction method [30] to obtain 8 B-scans, and each B-scan has 2048 A-lines across fast scan axis for 6.6 mm distance on retina. Each A-line averaged the 16 or 32 acquisition over ∼0.1mm distance in slow scan axis. Equivalently, the protocol averages 16 or 32 B-scans over 6.6 x 0.1mm slab. Therefore, the scanning protocol provided “noisy-clean” image pairs, where individual B-scans serve as noisy images and clean images are 16 or 32 averaging.

We have also manually segment 10 layers on the B-scan images, as shown in **Fig. 1**, each of which is individually reviewed. In order to reduce computation time, original data was down sampled four times to a size of 512×512, min-max normalized, and is stored as 16-bit TIFF files. In total, 105 noisy-clean image pairs were included. The dataset is from 12 normal subjects with varying image qualities.

**Fig. 1.**
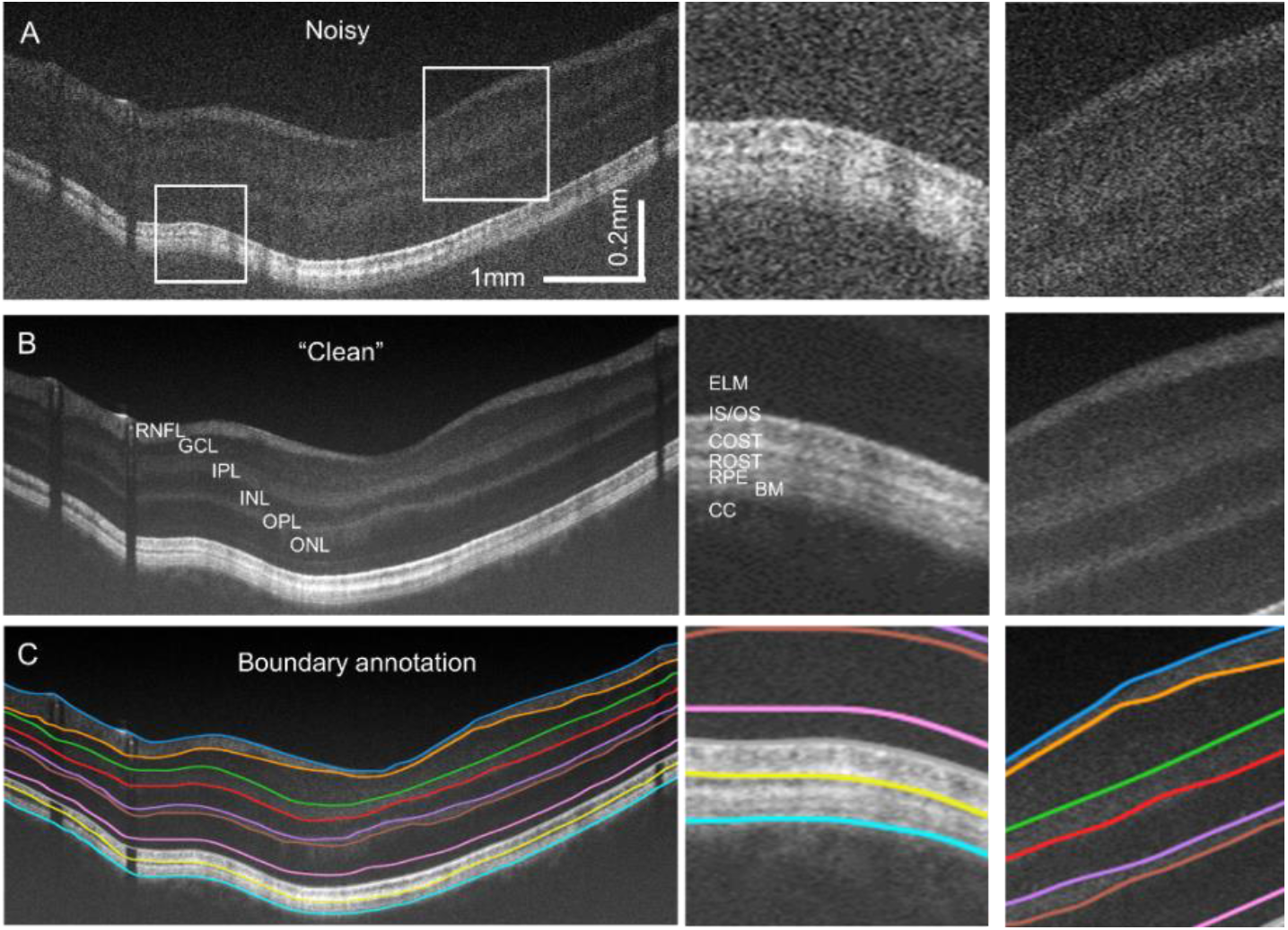
(A-B) A noisy-clean B-scan pair with (C) 10 manually delineated retinal boundaries. The anatomical layers are: retinal nerve fiber layer (RNFL); ganglion cell layer (GCL); inner plexiform layer (IPL); inner nuclear layer (INL); outer plexiform layer (OPL); outer nuclear layer (ONL); external limiting membrane (ELM); the inner segment (IS); the outer segment (OS); cone outer segment tip (COST); rod outer segment tip (ROST); retinal pigment epithelium (RPE); Bruch’s Membrane (BM) and choriocapillaris (CC). Zoom-in views from two small regions in inner and outer retina were displayed for comparison.

In order to benchmark the image quality of the dataset, we measured the mean value (Mean), standard deviation (Std) and contrast to noise ratio (CNR) of 11 regions in the images: the vitreous (the background region above RNFL, named UpperBg), 9 retinal layers and the choroid (the background region under BM, named LowerBg) and average the metrics over the dataset (**Table 1**). The contrast to noise ratio is calculated by:

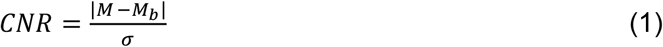

where M is the mean value of a certain layer *M*_*b*_ i s the mean value of the upper background and *σ* is the standard deviation of that region.

**Table 1.** Measured metrics of the VIS-OCT dataset.

|  | Mean (clean) | Mean (noisy) | Std (clean) | Std (noisy) | CNR (clean) | CNR (noisy) |
| --- | --- | --- | --- | --- | --- | --- |
| UpperBg | 858.1427 | 4389.4242 | 1399.9407 | 2757.4844 | 0.0000 | 0.0000 |
| RNFL | 14143.9416 | 15984.3259 | 5701.5912 | 6475.6281 | 3.2003 | 2.3298 |
| GCL | 9412.2750 | 11972.7236 | 3054.5016 | 4453.7112 | 3.6004 | 2.0473 |
| IPL | 11103.7092 | 13320.2133 | 3061.7223 | 4597.1574 | 4.3039 | 2.3560 |
| ICL | 9432.6936 | 11961.5811 | 2582.3825 | 4212.2997 | 4.1282 | 2.1270 |
| OPL | 12050.9172 | 14082.1736 | 3160.7442 | 4730.4772 | 4.5790 | 2.5034 |
| ONL | 9542.6530 | 12047.6997 | 2387.2608 | 4121.0761 | 4.4379 | 2.1842 |
| IS/OS | 11576.8291 | 13752.9136 | 2888.7658 | 4632.6602 | 4.7221 | 2.4562 |
| COST | 27473.6097 | 27902.3518 | 9276.7857 | 9971.7290 | 4.0120 | 3.2140 |
| ROST+RPE | 25916.0761 | 26428.4022 | 6906.3938 | 8090.3671 | 5.0288 | 3.6465 |
| LowerBg | 14217.2960 | 15971.9887 | 2973.5186 | 4799.9092 | 5.7484 | 2.9591 |

### 2.2. Network architecture

In the proposed DenoiSegOCT framework (**Fig. 2**), we utilize UNet-like encoder-decoder architecture for both denoising and segmentation tasks. The base block of each level of encoders and decoders includes 2 convolutional kernels with the residual operation to take advantage of residual learning [31] that improves the gradient flow during the optimization process. The depth = 5 is necessary for two reasons to *1)* provide large enough receptive field to capture the global information (i.e., the order of the retinal layers, which is the important prior anatomical knowledge in the task, and *2)* deep enough (i.e., more parameters and non-linear units) for the model to learn high-level features like the certain object of the layers and the order of the layers. The initial number of feature maps is 96. The input is the noisy 512x512 B-scan image. The training labels include manual segmentation boundaries, as well as labels for denoising. The denoising label depends on different strategies explained below.

**Fig. 2.**
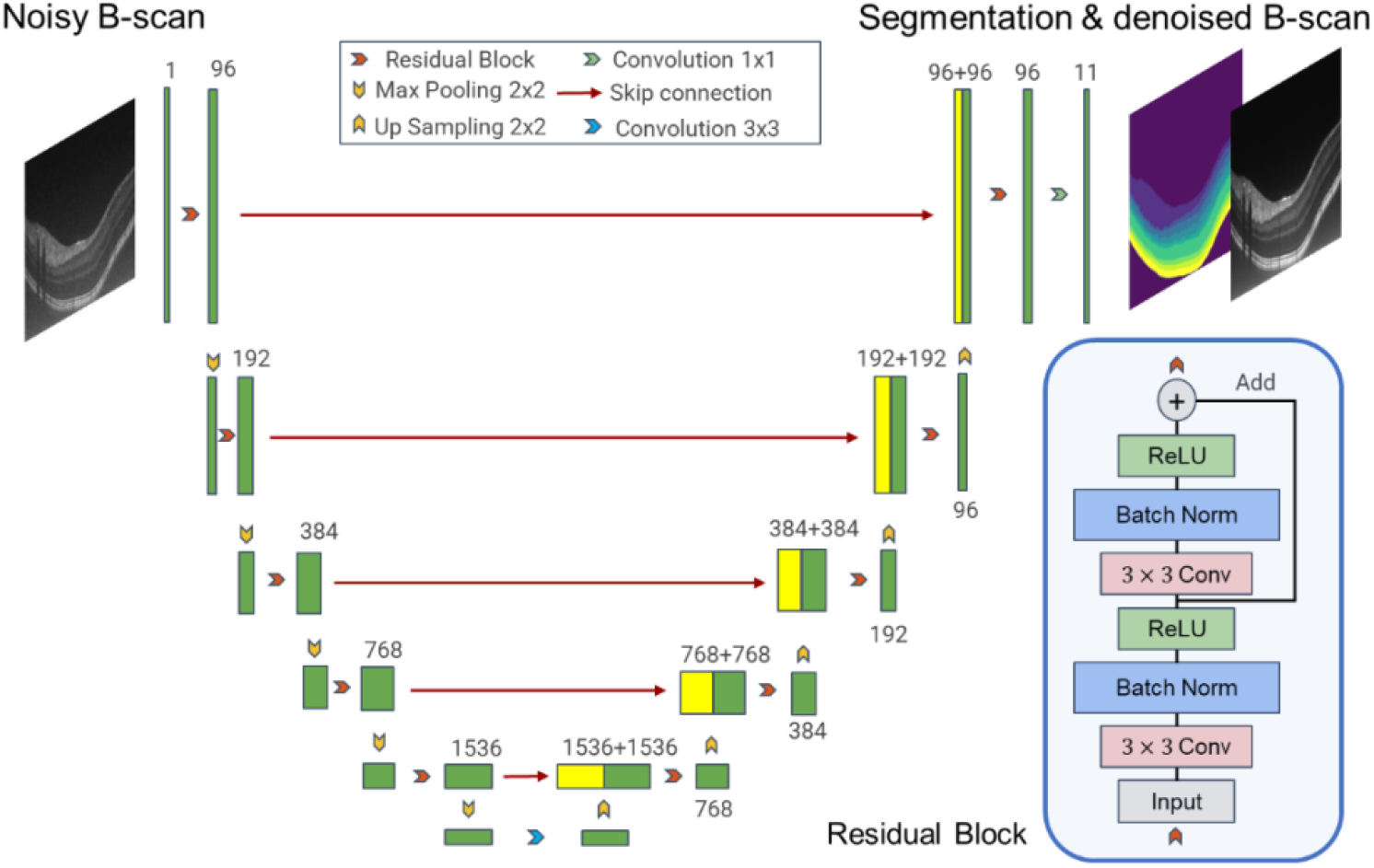
Architecture of DenoiSegOCT

### 2.3. Self-supervised denoising

We use the Noise2Void strategy (N2V) [28] to reduce the noises including the inherent speckle noise of VIS-OCT by adding an additional channel to the output layer of the network. N2V is a self-supervised denoising method that randomly selects and modifies several pixels (as blind spots of the network) in the image and then trains a neural network to restore the blind spots to original pixel values by optimizing mean squared error (MSE) loss of the two. During the training process, the network learns to predict the modified pixels by looking at a surrounding area, which is the receptive field of the network. We assume the noises in OCT are pixel-independent and the underlying content is pixel-dependent, thus this process can restore the content information degraded by the noises.

In this self-supervised denoising, the input is the modified image and the label is the original one. The modified pixels are substituted by a randomly selected pixel of their surrounding areas. In our study, 1.5 percent of the pixels in the input image were modified after the random batch cropping.

### 2.4. Supervised denoising

In the ideal case where clean ground truth is available, the “clean” B-scan ground truth is preferred as the label to supervise the training of the denoising model, named Noise2Label (N2L). We used MSE loss to optimize the network as in the N2V.

### 2.5. Ten-class segmentation

The pixel-wise ground truth mask, including 9 retinal layers and the background for the 10-class segmentation task, is created by filling pixels between the delineated retinal boundaries. We use a weighted cross-entropy loss with empirical weights for each class to alleviate the data-imbalance problem.

### 2.6. Co-learning strategy

The output layer of the network consists of 11 channels, of which one channel is the output of the denoised image and the remaining 10 channels provide the probability that each pixel belongs to the corresponding classes (*i*.*e*. retinal layers). The network is jointly trained by optimizing a combined loss with a task weight factor α and class weight factors *w*_*i*_:

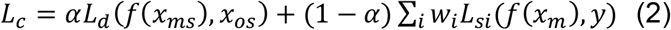

where *L*_*d*_ and *L*_*si*_ are the MSE loss for denoising and cross entropy loss for segmentation of class *i*; *f*(x) denotes the output of the network given an input *x*. The terms *x*_*ms*_, *x*_*os*_, and *x*_*m*_ denotes different images in three variations of DenoiSegOCT as following.

#### DenoiSegOCT(N2V)

The denoising component is self-supervised as described in Section 2.3. The *x*_*ms*_ and *x*_*os*_ are selected pixels of the modified and original images, respectively. *x*_*m*_ is the modified image and *y* is the corresponding pixel-wise segmentation label. The task weight factor α was set to 0.5 and class weight factors *w*_*i*_ were set to 0.5 for the background and 1 for the other classes.

#### DenoiSegOCT(N2L)

The denoising component is supervised by clean B-scans. The *x*_*ms*_ and *x*_*os*_ are replaced by the noisy image input and the clean B-scan label, respectively, and *x*_*m*_ is also replaced by the noisy image. The setting for α and *w*_*i*_ stays the same.

#### DenoiSegOCT(noisy)

The value of α is set to zero to ablate the denoising component in order to evaluate whether the co-learning strategy improves the performance of segmentation.

## 3. Experiments

### 3.1. Baselines and study comparison

To evaluate the performance of our DenoiSegOCT model, we first established baselines.

For segmentation, we selected the standard UNet architecture with 64 initial feature maps and a depth of 4 using either noisy or clean images as input [*i*.*e*. UNet (clean/noisy)]. In addition, we conducted experiments of sequential strategy that first denoised the image using a N2V model pretrained on our dataset and then trained the UNet segmentation (*i*.*e*. N2V+UNet). The pretrained N2V also serves as a baseline for self-supervised denoising.

For the proposed DenoiSegOCT, we embed both supervised denoising (N2L) and self-supervised denoising (N2V) into the framework, named DenoiSegOCT(N2L) and DenoiSegOCT(N2V). As for segmentation, the amount of annotated training data was set to 100%, 50%, and 25% to evaluate the label-efficient property of DenoiSegOCT. The loss of segmentation was set to zero for those images whose labels are unavailable. We finally set the weight factor α=0 in Equ. 2 to make it a pure segmentation task [DenoiSegOCT(noisy)] to evaluate the importance of the self-supervised denoising process when the amount of annotated training data is dropping.

### 3.2. Dataset, preprocessing and parameters

To the best of our knowledge, our VIS-OCT dataset is the first one available for simultaneous speckle noise reduction and segmentation. The dataset contains retinal images of 12 subjects (105 B-scans), of which 3 subjects (51 B-scans) were used for training, 4 subjects (17 B-scans) were used for validation and 5 subjects (37 B-scans) were used for test samples.

The network was implemented using Tensorflow and optimized by Adam optimizer. The training parameters are shown in **Table 2**, where the 8-fold augmentation included horizontal flipping and rotations. The initial learning rate was halved if the loss on the validation set did not decrease over ten epochs. For DenoiSegOCT(N2V) and N2V alone, 1.5 percent of the pixels (983 pixels as the blind spots in 2.3) in the input image were modified before the random batch cropping. The network was trained for 200 epochs and the model with the lowest validation loss was selected to test.

**Table 2.** Training parameters of baselines and DenoiSegOCT (Ours).

| Methods | Data augmentation | Learning rate | Input (crop) size | Batch size |
| --- | --- | --- | --- | --- |
| UNet (clean/noisy) | horizontal flipping | 0.0004 | 256×256 | 12 |
| N2V | 8-fold augmentation | 0.0004 | 64×64 | 128 |
| Ours (N2V) | horizontal flipping | 0.0004 | 256×256 | 12 |
| Ours (N2L) | horizontal flipping | 0.0002 | 512×512 | 2 |

## 4. Results

All the experiments were repeated 8 times and the mean and standard deviation are presented.

### 4.1. Visualization of the overall DenoiSegOCT performance

Figure 3 shows a representative example of the denoising and segmentation performance in DenoiSegOCT. For denoising, both N2V and DenoiSegOCT(N2V/N2L) significantly improved the image quality. Comparing N2V (Fig. 3I, J) to DenoiSegOCT(N2V) (Fig. 3M, N), higher contrast and shaper edges are observed using DenoiSegOCT than that using N2V alone (indicated by yellow arrows). In addition, the performance of DenoiSegOCT(N2L) (Fig. 3Q, R) has the best performance qualitatively and quantitively (**Table 3**).

**Fig. 3.**
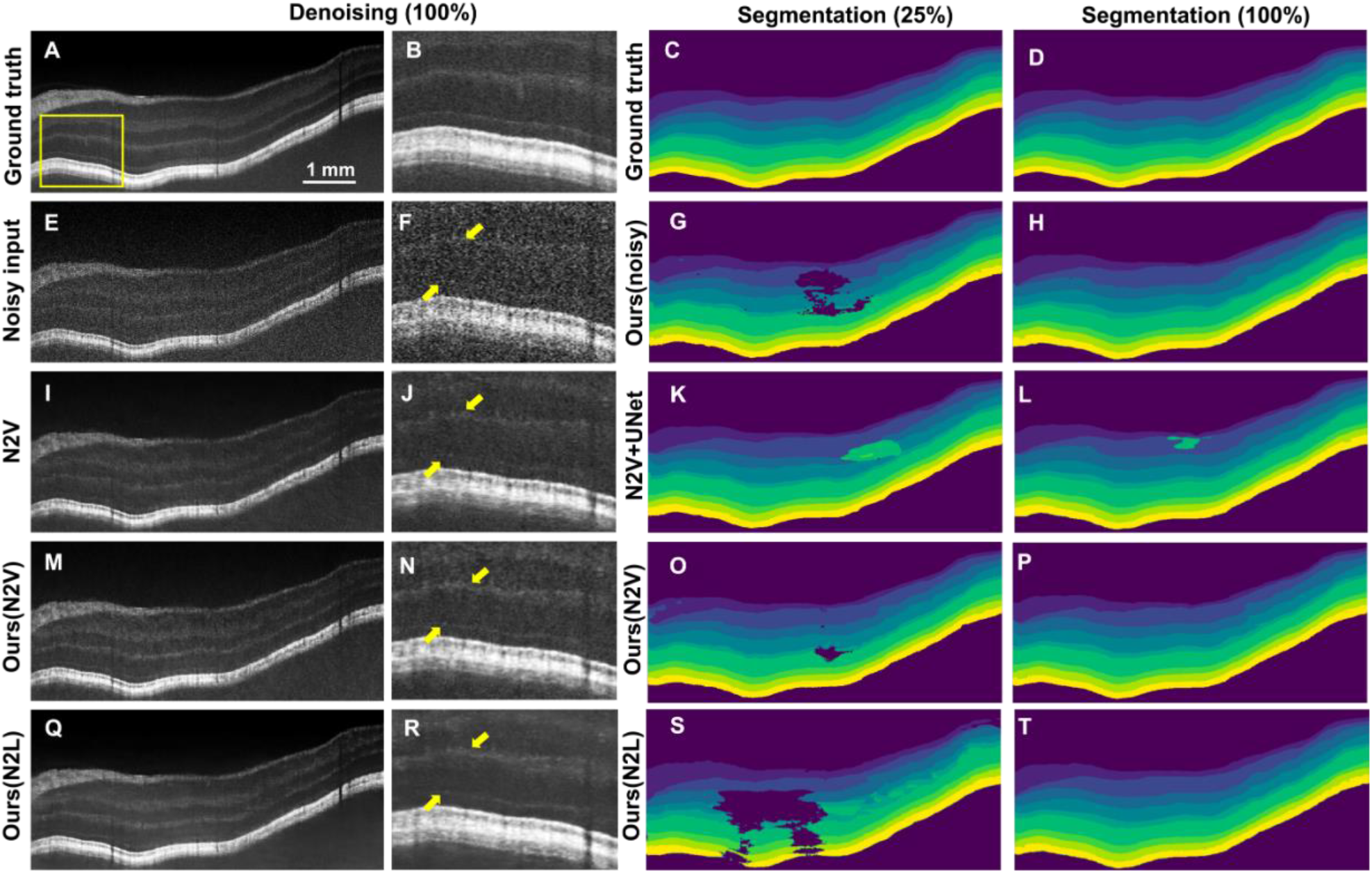
Overall denoising and segmentation visualization. A is the “clean” ground truth, E is the noisy input, I is the N2V denoised image, M is the DenoiSegOCT(N2V) denoised image, Q is the DenoiSegOCT (N2L) denoised image, and B, F, J, N, R are the corresponding zoom-in ROI shown as yellow box. The column of C, G, K, O, S is when 25% segmentation annotation is available, where C is the manual segmentation mask. G, K, O, S are the predicted segmentation masks using DenoiSegOCT without denoising, using Unet with image denoised by N2V, using ours with N2V denoising, and using ours with N2L denoising, respectively. The column of D, H, L, P, when 100% segmentation annotation is available. Bar = 1 mm.

**Table 3.** PSNR and SSIM of N2V denoised image baseline (N2V), self-supervised DenoiSegOCT [Ours(N2V) 100,50,25] with 100%, 50% and 25% segmentation annotation and supervised DenoiSegOCT [Ours(N2L) 100] with 100% segmentation annotation.

| Method | Noisy | N2V | Ours(N2V)100 | Ours(N2V)50 |
| --- | --- | --- | --- | --- |
| PSNR | 22.90 | 27.26($\pm 0.2470$) | 27.29($\pm 0.1503$) | 27.17($\pm 0.1552$) |
| SSIM | 0.2639 | 0.5296($\pm 0.0083$) | 0.5199($\pm 0.0054$) | 0.5180($\pm 0.0055$) |
| **Method** | ***Ours(N2V)25*** | ***Ours(N2L)100*** | ***Ours(N2L)50*** | ***Ours(N2L)25*** |
| PSNR | 26.96($\pm 0.2760$) | 30.20($\pm 0.1041$) | 29.07($\pm 0.0933$) | 27.35($\pm 0.6333$) |
| SSIM | 0.5145($\pm 0.0023$) | 0.7486($\pm 0.0056$) | 0.7024($\pm 0.0101$) | 0.6562($\pm 0.0147$) |

For segmentation with 100% annotation, all methods have excellent performance (Fig. 3 H, L, P, T). However, with only 25% annotation, comparing DenoiSegOCT(noisy) in Fig. 3G, DenoiSegOCT(N2V) (Fig. 3O) showed significantly improved performance for GCL, IPL and INL. We observe that the performance of DenoiSegOCT (N2L) degrades significantly in the case of only 25% annotations available (Fig. 3S).

### 4.2. Self-supervised and supervised denoising

We next quantitively evaluated our denoising performance using peak signal noise ratio (PSNR) and structure similarity index (SSIM) in Table 3. Using self-supervised strategy, both N2V and DenoiSegOCT(N2V) significantly increased the average PSNR/SSIM of noisy images. Note that the annotation amount for segmentation does not significantly affect the self-supervised denoising [*i*.*e*. Ours(N2V)100 vs. Ours(N2V)25 in Table 3]. While PSNR and SSIM in DenoiSeg(N2V)100 and N2V is similar, the denoised images by our framework is visually perceived with slightly sharper edges and higher contrast (Fig. 3 I, J and M, N), indicating segmentation may help denoising task in our co-learning network. In the supervised mode (*i*.*e*., N2L), the image quality was further boosted qualitatively and quantitively [Fig. 3 Q, R and Ours(N2L)100, 50 and 25 in Table 3].

### 4.3. Segmentation

Dice coefficient is used to evaluate the segmentation performance (**Table 4**). With 100% annotation available (Dice100), DenoiSegOCT(noisy) achieves comparable Dice coefficients (0.8727) with UNet(clean) (0.8852) and the sequential strategy N2V+UNet (0.8771); and is superior to UNet(noisy) (0.8710). In particular, DenoiSegOCT(N2V) becomes more advantageous as the number of available annotations decreases (Dice50 and Dice25 in Table 4), which indicates the synergetic enhancement of self-denoising to the layer segmentations.

**Table 4.**
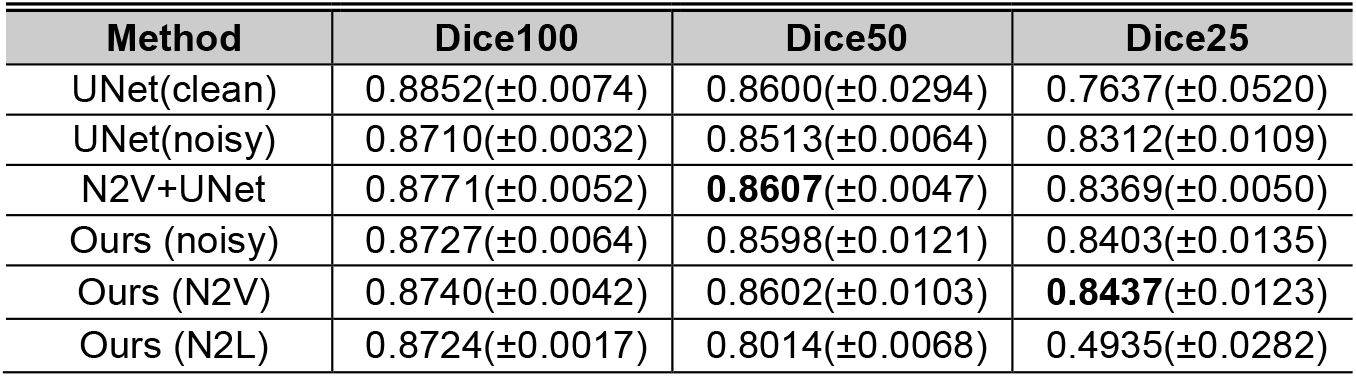
Dice coefficient for Unet baselines with clean input and noisy input, N2V+Unet, and our DenoiSegOCT(noisy), DenoiSegOCT(N2V) and DenoiSegOCT (N2L). Ours is in short for DenoiSegOCT. Dice100, Dice50, and Dice25 represent 100%, 50%, 25% of the annotation are available, respectively.

We further conducted an ablation study in **Table 5** that comparing our model with and without N2V denoising process in the segmentation task. When available annotation drops to 25%, significant improvement (∼2% higher Dice over all test data and all 8 experiments) is found in DenoiSegOCT(N2V) over DenoiSegOCT(noisy) for GCL, IPL and INL. These layers are the most blur and ambiguous regions in the B-scan because of the low signal to noise ratio. At those regions, the features (*e*.*g*., edges) are difficult to distinguish and segment for both naked eyes and machine. This comparison suggests the DenoiSegOCT(N2V) efficiently performs both segmentation and self-denoising simultaneously when label annotation is limited.

**Table 5.**
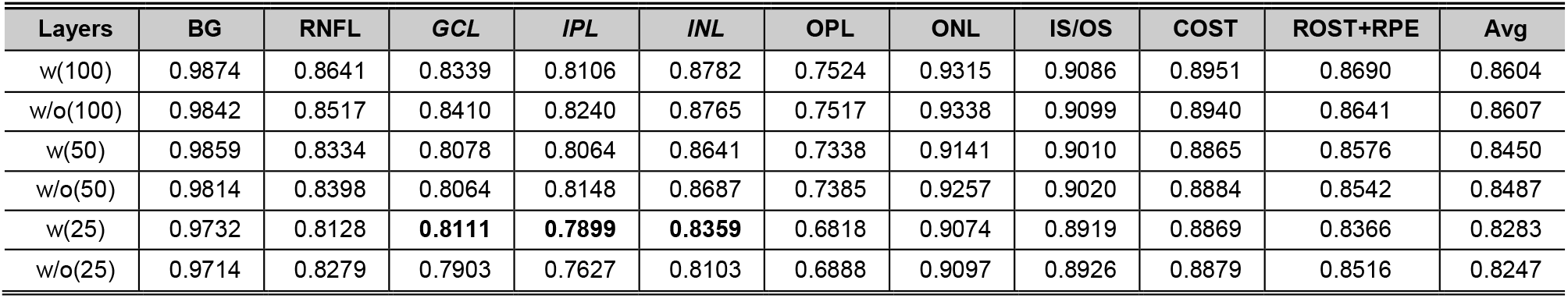
Dice coefficient of background (BG) and all 9 retinal layers with(w) or without(w/o) N2V denoising process using DenoiSegOCT when 100, 50, 25% segmentation annotation is available.

| Layers | BG | RNFL | GCL | IPL | INL | OPL | ONL | IS/OS | COST | ROST+RPE | Avg |
| --- | --- | --- | --- | --- | --- | --- | --- | --- | --- | --- | --- |
| w(100) | 0.9874 | 0.8641 | 0.8339 | 0.8106 | 0.8782 | 0.7524 | 0.9315 | 0.9086 | 0.8951 | 0.8690 | 0.8604 |
| w/o(100) | 0.9842 | 0.8517 | 0.8410 | 0.8240 | 0.8765 | 0.7517 | 0.9338 | 0.9099 | 0.8940 | 0.8641 | 0.8607 |
| w(50) | 0.9859 | 0.8334 | 0.8078 | 0.8064 | 0.8641 | 0.7338 | 0.9141 | 0.9010 | 0.8865 | 0.8576 | 0.8450 |
| w/o(50) | 0.9814 | 0.8398 | 0.8064 | 0.8148 | 0.8687 | 0.7385 | 0.9257 | 0.9020 | 0.8884 | 0.8542 | 0.8487 |
| w(25) | 0.9732 | 0.8128 | **0.8111** | **0.7899** | **0.8359** | 0.6818 | 0.9074 | 0.8919 | 0.8869 | 0.8366 | 0.8283 |
| w/o(25) | 0.9714 | 0.8279 | 0.7903 | 0.7627 | 0.8103 | 0.6888 | 0.9097 | 0.8926 | 0.8879 | 0.8516 | 0.8247 |

Counterintuitively, the segmentation performance of DenoiSegOCT(N2L) degrades significantly as the number of available annotations decreases, from 0.8724 to 0.4935 in **Table 4**.

### 4.4. Generalization ability

We continued to test whether the model trained on our proposed HD dataset can generalize well to the raster scanning dataset, which was obtained from a totally different scanning protocol using our device, thereby denoise and segment the whole 3D volume (**Fig. 4**). The range of raster protocol was 3 mm × 3 mm with 512 A-lines × 512 B-scan across the retina. The raster protocol does not include A-line modulation like HD protocol so only noisy images are available. The noisy B-scan images followed the same preprocessing steps as training data as described in Section 2.1.

**Fig. 4.**
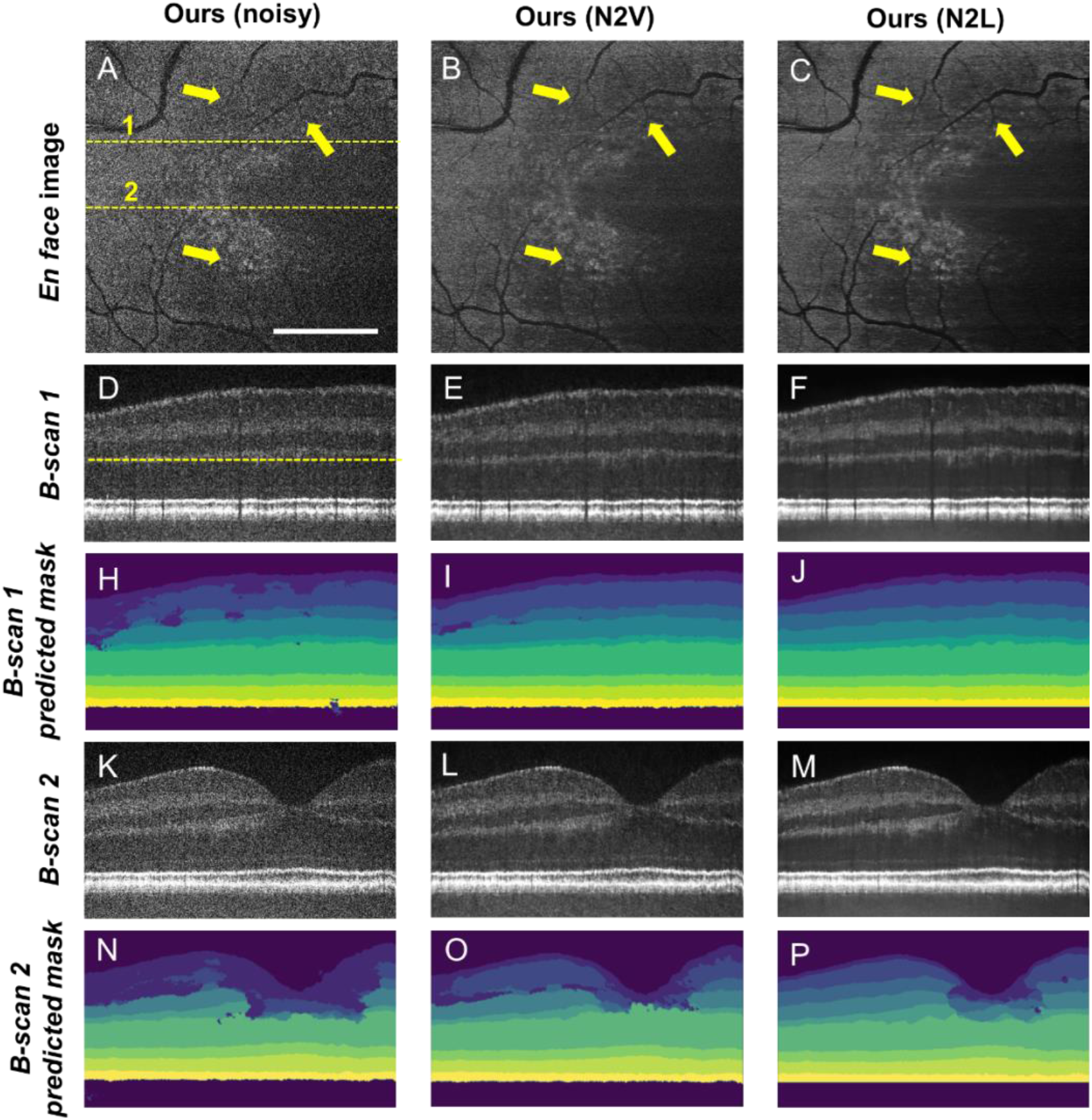
Visualization of denoising and segmentation on a raster scanning 3D dataset. A, D, J are from the noisy input volume, B, E, K/C, F, L are from the volumes denoised by DenoiSegOCT with N2V/N2L. G, H, I are predicted segmentation masks for B-scan 1 from DenoiSegOCT without denoising process, DenoiSegOCT with N2V and DenoiSegOCT with N2L, respectively. M, N, O are predicted segmentation masks for B-scan 2. B-scan 1 and B-scan 2 are from cross section 1 and 2 (yellow dashed lines in A). *En face* images A, B and C are one depth section around OPL (yellow dashed lines in D). The retina was flattened at RPE for better visualization of the layer segmentation. Bar= 1 mm.

Figure 5A-5C compares the original image and two of our proposed models at one depth section around OPL. The noise has been effectively suppressed to enhance features such as small vessel branches (Fig. 5B, C). Two representative B-scans, one is from the perifovea (B-scan 1) and another crosses the fovea pit (B-scan 2), demonstrate the denoising and segmentation performance of our models. Qualitatively, both DenoiSegOCT(N2V) and DenoiSegOCT(N2L) increase the image quality significantly while DenoiSegOCT(N2L) is better (Figure. 4 D, E, F, J, K, L).

For segmentation, we observed that the denoising process helped the segmentation by comparing Fig. 4G, H and I, as well as Fig. 4 M, N and O, while DenoiSegOCT(N2L) performs the best.

## 5. Discussion

In the paper, we presented the first VIS-OCT dataset from normal eyes with noisy-clean pairs to the scientific community. We investigated a DL framework [DenoiSegOCT(N2V)] to simultaneously perform self-denoising and segmentation which in turn synergistically improved both tasks within the same network. The two-in-one design also shows improved efficiency when the segmentation annotation is scarce (25%) in some difficult layers, such as GCL, IPL, and INL.

In addition, we compared the denoising performance with self-supervised (N2V) and supervised (N2L) model in DenoiSegOCT. It is not surprising that supervised model with “clean” ground truth image performs better. Also, the assumption of N2V that the noises in the image are pixel-independent may not be strictly held (e.g., the speckle noise) which may impact the self-denoising. Future work including speckle statistics may further improve the performance in N2V. While N2L outperformed N2V in DenoiSegOCT, it is counterintuitive that segmentation performance in DenoiSegOCT (N2L) degrades when annotated segmentation labels are reduced (Table 4, last row). We speculate the following reasons. 1) The N2L is a mapping from the noisy image to the clean image, from where the neural network learns inter-relationship of the two images. On the other hand, in the N2V the neural network learns intra-relationship of the noisy image itself, indicating the N2V can help to learn the self-features of the noisy image, e.g., the correlation of retinal layers while N2L cannot. 2) The learning process of N2L is simpler than N2V, thus the gradient drops much faster during training. On the other hand, the gradient of segmentation branch is not enough to update the parameters when the segmentation labels are not enough. Therefore, the neural network converged quickly to a local minimum, became “lazy” when the denoising branch was trained sufficiently.

It is interesting to observe the segmentation was improved in difficult layers (GCL, INL, IPL) with the help of self-denoising when the layer annotation reduced on 25%. Normally the supervision of annotation can guide the model to distinguish the blur regions in the B-scans if there is a fair amount of training data. However, we believe that when the annotation amount decreases significantly, the N2V denoising process takes over and helps capture similar and useful features for segmentation in DenoiSegOCT. One exception is for OPL, which is the most challenging because it is thin and irregular, resulting unstable training with few annotations. This observation aligns with a previous report [21]. The synergetic improvement on segmentation is also evident in Fig. 4. Given the same noisy input without “clean” ground truth, the DenoiSegOCT(N2V) performed better in layer segmentation than without self-denoising (Fig. 4G vs 4H, Fig. 4M vs 4N).

The trained DenoiSegOCT generalized reasonably well on images taken with the raster scanning protocol (Fig. 4), particularly on the denoising task. We noted that the segmentation error is more prominent in the B-scans crossing the fovea pit (Fig. 4M-4O). The reason is likely that 1) there is less training data on B-scans crossing the fovea pit than other retinal sections, and 2) several inner retinal layers converge and disappeared at the fovea pit which are more challenging to segment than layers with uniform thickness.

## 6. Conclusion

In this paper, we present the first VIS-OCT retinal image dataset for data-driven method development. The co-learning framework DenoiSegOCT is efficient to simultaneously denoise and segment retinal layers. The synergy between two tasks within the same network improves each other, particularly with segmentation annotation is limited. The models trained on our proposed dataset generalized well to the unseen raster scanning dataset, indicating the robustness of our framework.

## Acknowledgements

This study was supported by NIH R01NS108464, and R01EY032163.

## Data availability

The complete VIS-OCT dataset will be made public on JHU data repository upon acceptance of the manuscript.

